# Age- and sex-related differences in baboon (*Papio anubis*) gray matter covariation

**DOI:** 10.1101/2021.12.08.471755

**Authors:** M. M. Mulholland, A. Meguerditchian, W. D. Hopkins

## Abstract

Age-related changes in cognition, brain morphology, and behavior are exhibited in several primate species. Baboons, like humans, naturally develop Alzheimer’s disease-like pathology and cognitive declines with age and are an underutilized model for studies of aging. To determine age-related differences in gray matter covariation of 89 olive baboons *(Papio anubis*), we used source-based morphometry (SBM) to analyze data from magnetic resonance images. We hypothesized that we would find significant age effects in one or more SBM components, particularly those which include regions influenced by age in humans and other nonhuman primates (NHPs). A multivariate analysis of variance revealed that individual weighted gray matter covariation scores differed across the age classes. Elderly baboons contributed significantly less to gray matter covariation components including the brainstem, superior parietal cortex, thalamus, and pallidum compared to juveniles, and middle and superior frontal cortex compared to juveniles and young adults (*p*<0.05). Future studies should examine the relationship between the changes in gray matter covariation reported here and age-related cognitive decline.

## 1 Introduction

Both humans and nonhuman primates show changes in cognition, brain morphology, and behavior with age, both as a part of normal aging and as a result of age-related disease. Normal human aging is characterized by declines in cognition, global brain volume, gray matter volume, and functional activation associated with increasing age (Abe et al., 2008; Bigler et al., 2002; Bigler et al., 1997; Bishop et al., 2010; Brickman et al., 2007; Chen et al., 2013; Ge et al., 2002; Good et al., 2001; Hawkins et al., 2015). Most studies on nonhuman aging have utilized non-primate model species such as yeast, nematodes, fruit flies, and mice due mostly to availability, cost, short lifespan and well-documented genetic and/or physiological characteristics. While these models have provided important data and have advanced our understanding of aging, they do not closely model the human aging process. Utilizing nonhuman primate models (particularly those more closely related phylogenetically) is critically important given their genetic similarities to humans, longer lifespans, larger and more complex brains and more sophisticated cognitive abilities compared to other model species. Among nonhuman primates, baboons are particularly well-suited as a model for human aging. First, unlike other monkey species, there is evidence that baboons undergo well-coordinated growth, growth spurts, and an order of growth cessation like humans (Leigh, 2009). Second, the baboon genome is very similar to humans with over 90% homology of coding segments and 85% homology of non-coding regions. For this reason, the baboon has served as a model to study the genetics and epigenetics of several human diseases (Cox et al., 2013; Cox et al., 2006; Robinson et al., 2019; Rogers et al., 2009; Rogers et al., 2000). In addition, both human and baboon lifespan is moderately heritable (0.23-0.26 and 0.23, respectively) and differences in frailty likely causes variation in individual mortality risk within both species (Bronikowski et al., 2002; Martin et al., 2002). Third, baboons have been shown to develop similar age-related diseases and conditions as humans, including osteoporosis and arthritis, menopause, endometriosis, obesity, diabetes, cardiovascular disease, and Alzheimer’s-like pathology (Martin et al., 2002; Schultz et al., 2000a; Schultz et al., 2000b; Schultz et al., 2001; Yeung et al., 2016). For example, postmortem studies have shown that baboons develop tau pathology in both neurons and glia and Aβ plaques as they age (Schultz et al., 2000a; Schultz et al., 2002; Schultz et al., 2000b; Schultz et al., 2001). Furthermore, compared to rhesus macaques (a common nonhuman primate model species), baboons have larger brains making them ideal for imaging studies (Black et al., 2009), and the baboon prefrontal cortex is more developed and more similar to the human prefrontal cortex than the rhesus macaque (Brent, 2009; Fridman and Popova, 1988). Due to these characteristics, studies of baboon aging can help bridge the translational gap between other model species and human aging studies.

Like humans, there is evidence of age-related cognitive decline in baboons. In a cross-sectional study of 19 baboons (aged 1-14), Bonté (2014) found that executive control (as measured by a transfer index task) declined with age. In another cross-sectional study of 6 baboons, older adults (20-23 years old) performed significantly worse on cognitive tasks (specifically, learning a novel task, precision, and simple discrimination) compared to younger adults (13-16 years old) (Lizarraga et al., 2020). These findings suggest that there are likely neurobiological changes occurring as baboons age. Although baboon neuroanatomy and function is well-studied (e.g., (Amiez et al., 2019; Atkinson et al., 2015; Black et al., 2009; Blaizot et al., 2004; Cain and Wada, 1979; Greer et al., 2002; Love et al., 2016; Marie et al., 2018; McBride et al., 1999; Meguerditchian et al., 2021; Rogers et al., 2007; Westerhausen and Meguerditchian, 2021), and postmortem studies reveal the development of tau pathology as they age (Schultz et al., 2000a; Schultz et al., 2002; Schultz et al., 2000b; Schultz et al., 2001), to date there have been few *in vivo* neuroimaging studies focused specifically on the aging baboon brain. Franke et al. (Franke et al., 2017) found that prenatal undernutrition (70% of normal daily food intake) had long-term effects on the baboon brain, prematurely aging the young adult female baboon brain by 2.7 years on average. In addition, the authors examined typical brain aging across the baboon lifespan and reported a significant decrease in overall gray matter volume and increase in overall white matter volume with age. More recently, Westerhausen and Meguerditchian (2021) examined corpus callosum morphology across the lifespan and found that the area of the corpus collosum increases slowly with age, particularly within the anterior section.

Here we examine the relationship between age and whole-brain gray matter covariation using source-based morphometry (SBM). SBM is a multivariate approach which incorporates an independent components analysis to create networks of brain regions that covary among gray or white matter voxels across the whole brain (Gupta et al., 2019). Here we examine the contribution of both age and sex to gray matter covariation within the baboon brain. A number of studies in human subjects have reported age-related changes in gray matter covariation networks (DuPre and Spreng, 2017; Evans, 2013; Hafkemeijer et al., 2014; Koini et al., 2018; Liu et al., 2017; Montembeault et al., 2012; Xu et al., 2009b), but there are only a few studies of this type in nonhuman primates. Source-based morphometry-like methods were used to show that older rhesus macaques had reduced gray matter covariation in components including dorsal and ventral prefrontal cortex, superior temporal sulcus, and sylvian fissure (Alexander et al., 2008). Previous studies in chimpanzees have also reported age-related changes in gray matter covariation, with decreased gray matter covariation reported for the superior frontal, supplementary motor, and anterior temporal cortex as chimpanzees age (Hopkins, W. D. et al., 2019). In addition, they reported increased gray matter covariation in the prefrontal and premotor cortices and part of the cerebellum as chimpanzees age (Hopkins, W. D. et al., 2019). A more recent study of age-related changes in chimpanzee gray matter (using voxel-based morphometry) found significant reductions in the prefrontal cortex, middle temporal cortex, superior frontal cortex, insula, superior temporal cortex, and entorhinal cortex with increased age (Mulholland et al., 2021). In the current study, we use source-based morphometry to examine the relationship between age and gray matter covariation in a baboon sample. We hypothesized that significant age effects would be found in one or more of the SBM components, specifically those comprised of regions known to be influenced by age in humans and other non-human primates, such as the middle temporal lobe, prefrontal cortex, and the cerebellum.

In addition, we examined sex differences in gray matter covariation. Studies of the baboon brain have indicated that sex may account for variation in overall brain volume and sulcal length (Atkinson et al., 2015; Rogers et al., 2007), differences in the area of the corpus collosum (Phillips and Kochunov, 2011), and whole brain gray and white matter volumes (Franke et al., 2017). Franke et al. (2017) found that males had significantly higher absolute gray matter volume than females but did not examine regional differences or gray matter covariation. Atkinson et al. (2015) reported that sex accounted for a significant proportion of variance in the length of the central sulcus, lateral fissure, lunate, and superior temporal sulcus. In chimpanzees, an SBM analysis revealed significant sex differences in gray matter covariation, including the cerebellum, primary visual, anterior temporal, and frontopolar cortices (Hopkins, W. D. et al., 2019). Similar to chimpanzees, here we hypothesized that there would be significant sex differences in one or more of the baboon SBM components.

## 2 Materials and Methods

### 2.1 Subjects

We utilized MRI data collected from 89 captive olive baboons (56 females, 31 males) from the Station de Primatologie CNRS in Rousset, France (Agreement number for conducting experiments on vertebrate animals: D13-087-7). The baboons ranged in age from 2 to 26 years of age (*M* = 11.46, *SD* = 5.89). All subjects were socially housed with free access to indoor/outdoor enclosures furnished with wooden platforms and vertical climbing structures. Care staff fed the baboons a diet of commercial primate chow, seed, and fresh produce four times per day. All baboons had *ad libitum* access to water. All procedures complied with the current French laws and the European directive 86/609/CEE and were approved by the Provence Alpes Côte d’Azur ethics committee.

### 2.2 Image Acquisition and Post-Image Processing

Magnetic resonance image (MRI) scans were acquired using a 3.0 Tesla scanner (MEDSPEC 30/80 ADVANCE; Bruker) with a Rapid-Biomed surface antenna located at the Marseille Functional MRI Center. Detailed information about both image acquisition and post-image processing can be found in (Love et al., 2016). Briefly, each baboon was sedated with an intramuscular injection of ketamine (10 mg/kg) prior to transportation to the imaging facility. Upon arrival, subjects were further sedated with intramuscular injections of tiletamine and zolazepam (Zoletil™, 7 mg/kg) and acepromazine (Calmivet™, 0.2–0.5 mg/kg). During the imaging procedures, subjects were anesthesized with a drip irrigation of tiletamine, zolazepam (Zoletil™, 4 mg/kg/h) and NaCl (0.9% of 4 ml/kg/h). Veterinary staff monitored cardiovascular and respiratory functions with a SpO2 device and respiratory belt. Animals were placed into the scanner in a prone position with their head supported by cushions and Velcro strips. High resolution structural T1-weighted images were acquired (TR: 9.4 ms; TE: 4.3 ms; flip angle: 30°; inversion time: 800 ms); due to size field of view and isotropic voxel size differed for female and young male baboons (fov: 108 × 108 × 108 mm; isotropic voxel size: 0.6 mm3) and adult males (fov: 126 × 126 × 126 mm; isotropic voxel size: 0.7 mm3). Upon completion of the scan, subjects were monitored and fully recovered from anesthesia prior to being reunited with their social groups.

Resulting images were first denoised using an Adaptive Nonlocal Means filter (ANLM; (Manjón et al., 2010)) before brain extraction using both Multi Atlas Skull Stripping software (http://www.cbica.upenn.edu/sbia/software/MASS/index.html; (Doshi et al., 2013; Ou et al., 2011) and ITK-SNAP (version 3.2, www.itksnap.org, (Yushkevich et al., 2006). Next, the skull stripped images were bias corrected using an N4 algorithm (Tustison et al., 2010). Finally, we ran these images through the FSL-VBM pipeline using the FMRIB Software library tools (FSL; Oxford; http://fsl.fmrib.ox.ac.uk/fsl/fslwiki/FSLVBM). This process included (1) segmentation of each scan into gray and white matter, (2) linear registration of each scan to a standard baboon template (Love et al., 2016), (3) creation of a study-specific gray matter template (Andersson et al., 2007; Douaud et al., 2007; Smith et al., 2004), (4) non-linear registration of each subject’s gray matter image to the study-specific template, (5) modulation of the gray matter volume by use of a Jacobian warp to correct for local expansion or contraction of gray matter within each voxel, and (6) smoothing with an isotropic Gaussian kernel with a sigma of 2 mm. The resulting smooth, modulated gray matter volumes were used in subsequent analyses.

### 2.3 Statistical Analyses

#### Source-based morphometry

The data were analyzed using source-based morphometry (SBM v 1.0b) performed in the Group ICA of fMRI Toolbox (GIFT; http://icatb.sourceforge.net)(Xu et al., 2009a) in MATLAB R2015b. We imported the smoothed, modulated gray matter volumes for each subject and allowed the software to estimate the number of components based on an independent component analysis using a neural network algorithm. After estimation, the program calculates the parameters and creates a 4D volume containing each of the independent spatial components. At the individual level, SBM produces a weighted score that reflects each subject’s relative contribution to the creation of each component, which were then used in subsequent statistical analyses. To examine the brain regions contributing to each component, we registered the component maps (scaled to standard deviation units and z scores) to the standard baboon template brain (Love et al., 2016) and set the z-score threshold at |*z*| ≥ 3.00 as has been used in previous studies in humans and chimpanzees (see Hopkins, W. D. et al., 2019; Hopkins et al., 2020b; Xu et al., 2009a). All brain regions reaching this threshold were considered significant, and the volume was then measured using the region-of-interest tool in ANALYZE 14.

#### Age and sex effects

We ran a multivariate analysis of variance (MANOVA) with age group and sex (female, male) as the independent variables and weighted scores for each SBM component as dependent variables. Subsequent univariate ANOVAs were then used to examine differences between the age groups on their contribution to each individual SBM component.

## 3 Results

The SBM analysis identified 10 independent gray matter covariation components and corresponding individual weighted scores. The largest regions within each component included 1. cerebellum (bilateral), 2. caudate (bilateral) and putamen (bilateral), 3. brainstem (bilateral) and superior parietal cortex (bilateral), 4. isthmus of the cingulate (bilateral), 5. occipital cortex (bilateral), 6. brainstem (bilateral), 7. thalamus (bilateral), 8. middle frontal and superior frontal cortex (bilateral), 9. amygdala and hippocampus (bilateral), and 10. brainstem and cerebellum (bilateral). Full anatomical descriptions of each component and their respective gray matter volumes can be found in Table 1. 3D renderings of each component can be found in the supplemental materials.

**Table 1.** Regions and corresponding volumes (≥ 20 *mm*^*3*^) of the 10 baboon SBM components.

| Component | Region | Volume | Left/Right Volume |
| --- | --- | --- | --- |
| C1 | cerebellum | 3210.84 | 1678.75 / 1532.09 |
| C2 | putamen | 391.18 | 231.34 / 159.84 |
|  | cerebellum | 368.07 | 241.06 / 127.01 |
|  | caudate | 300.24 | 130.03 / 170.21 |
|  | lingual gyrus | 217.3 | 131.33 / 85.97 |
|  | inferior temporal cortex | 47.95 | 47.95 / 0 |
|  | brainstem | 43.85 | 43.85 / 0 |
| C3 | brainstem | 478.88 | 299.81 / 179.07 |
|  | superior parietal | 454.89 | 284.47 / 170.42 |
|  | thalamus & pallidum | 424.01 | 2.81 / 421.20 |
|  | precuneus | 53.35 | 44.28 / 9.07 |
|  | cerebellum | 39.53 | 15.77 / 23.82 |
| C4 | isthmus cingulate | 203.26 | 92.88 / 110.38 |
|  | insula | 98.71 | 0 / 98.71 |
|  | amygdala | 66.74 | 0 / 66.74 |
|  | inferior parietal | 63.72 | 39.96 / 23.76 |
|  | superior temporal cortex | 45.15 | 0 / 45.15 |
| C5 | occipital cortex | 3843.08 | 2098.66 / 1744.42 |
| C6 | brainstem | 1877.68 | 1010.66 / 867.02 |
| C7 | thalamus | 1879.2 | 909.58 / 969.62 |
|  | putamen & insula | 665.5 | 484.06 / 181.44 |
| C8 | middle frontal cortex | 1227.52 | 578.66 / 648.86 |
|  | superior frontal cortex | 1104.63 | 559.01 / 545.62 |
|  | precentral cortex and insula | 92.23 | 0 / 92.23 |
| C9 | amygdala & hippocampus | 2164.97 | 1295.57 / 869.40 |
|  | superior temporal cortex | 518.19 | 309.53 / 208.66 |
|  | rostral middle frontal cortex | 56.81 | 56.81 / 0 |
|  | pericalcarine | 49.68 | 49.68 / 0 |
|  | precuneus | 48.6 | 4.54 / 44.06 |
|  | brainstem | 28.29 | 11.66 / 16.63 |
| C10 | brainstem & cerebellum | 3570.05 | 2022.41 / 1547.64 |

Baboons were grouped by age: juvenile (2-6 years old), young adult (7-12 years old), middle age (13-16 years old), and geriatric (17+ years old). The MANOVA revealed significant main effects for age group [*F*(30, 222) = 3.245, *p* < 0.001 *η*^*2*^ = 0.305], and sex [*F*(10, 72) = 6.213, *p* < .001, *η*^*2*^ = 0.463], but no significant interaction between the two variables *F*(30, 222)=1.161, *p* > 0.05, *η*^*2*^=0.136. For age, subsequent univariate tests revealed significant effects for Component 3 *F*(3, 81) = 4.149, *p* < 0.01, Component 5 *F*(3, 81) = 6.100, *p* < 0.01, Component 7 *F*(3, 81) = 3.929, *p* < 0.02, and Component 8 *F*(3, 81) = 20.701, *p* < 0.001. The mean weighted scores for each age group on components 3, 5, 7, and 8 are shown in Figure 1. For Components 3 and 8, post-hoc analyses indicated that the weighted contribution to the components were significantly lower in elderly baboons compared to other age classes. Elderly baboons contributed significantly less to component 3 compared to juveniles, and less to component 8 than both juveniles and young adults. In contrast, for Component 5, elderly baboons contributed significantly more compared to juvenile baboons and for Component 7, elderly baboons contributed significantly more compared to both juvenile and young adult baboons.

**Figure 1.**
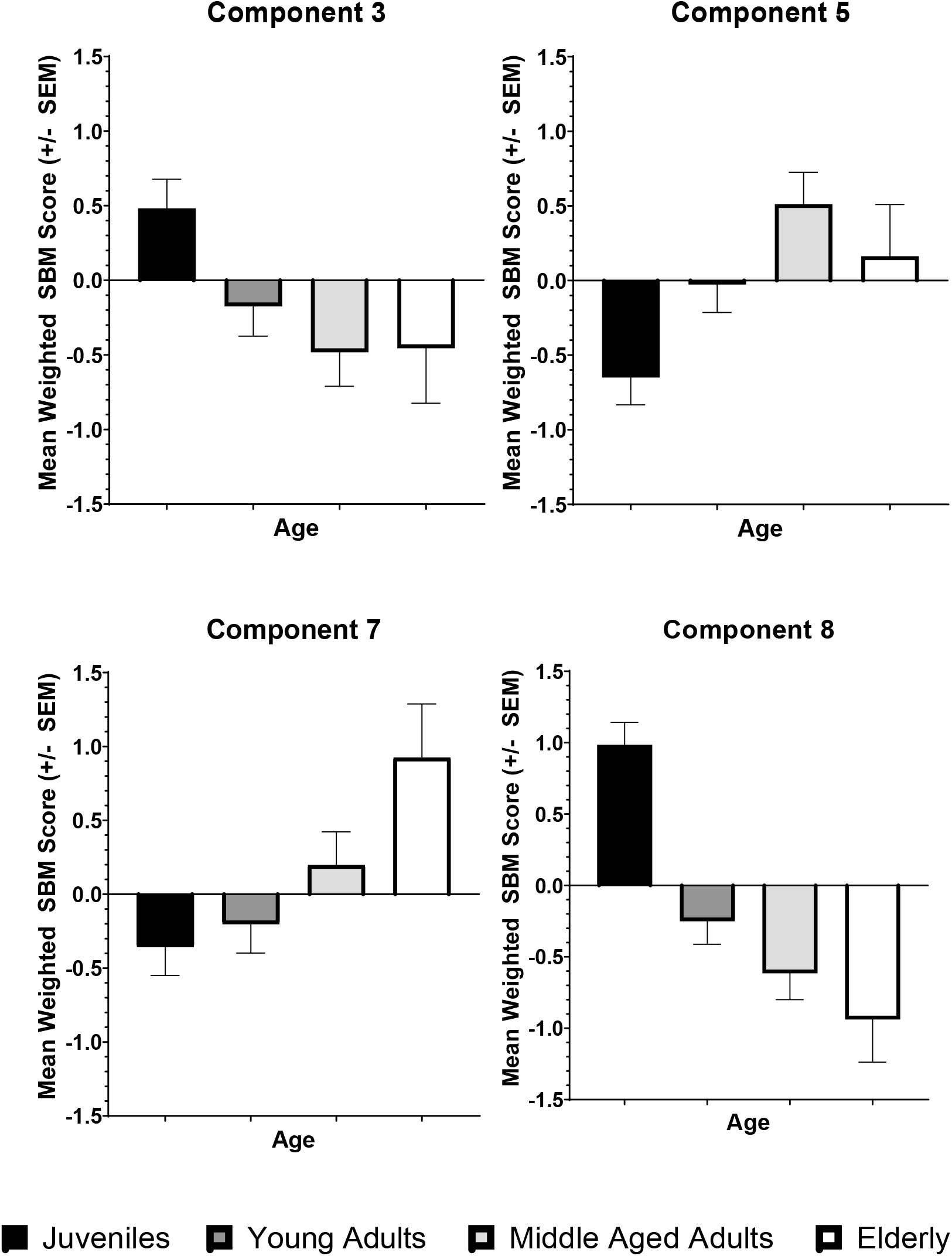
The average weighted SBM score on Components 3, 5, 7, and 8 for each age class. Elderly baboons contributed more to Components 5 and 7 but contributed less to components 3 and 8 compared to juvenile baboons (*p* < 0.05).

Regarding sex, univariate analyses revealed significant differences between males and females for Component 2 *F*(1, 81) = 14.068, *p* < 0.001, Component 3 *F*(1, 81) = 6.298, *p* < 0.05, Component 8 *F*(1, 81) = 5.398, *p* < 0.05, and Component 9 *F*(1, 81) = 4.819, *p* < 0.05. For Components 3, 8, and 9, males contributed less to the component scores than females, whereas females contributed less to component 2 compared to males. The mean weighted scores for each sex and component are shown in Figure 2.

**Figure 2.**
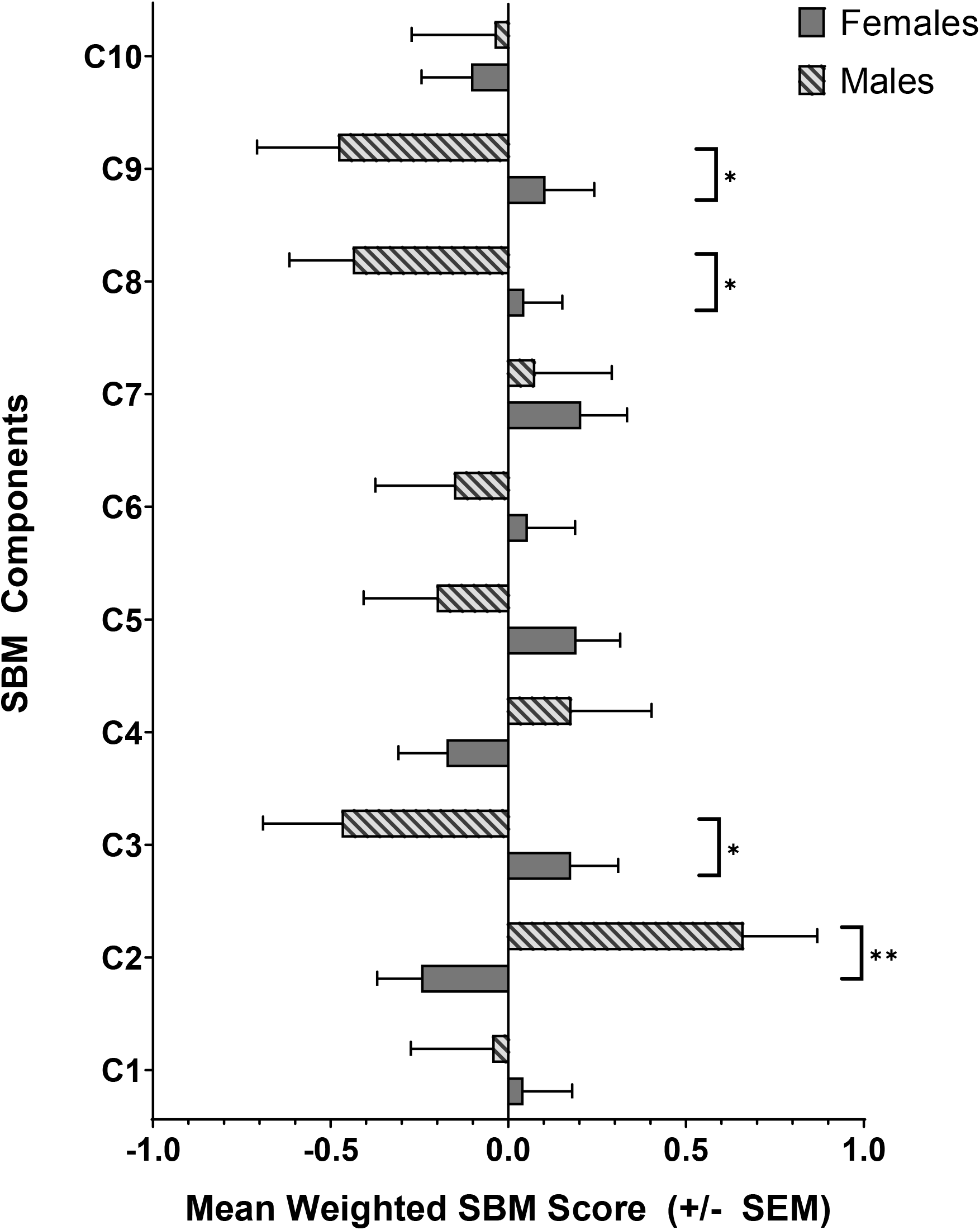
The average weighted SBM score on each SBM component for males and females. Males contributed more to component 2, whereas females contributed more to components 3, 8, and 9. * *p* < 0.05, ** *p* < 0.01

## 4 Discussion

Using source-based morphometry, we identified 10 components of gray matter covariation in the baboon brain and examined the relationship with age. First, contribution to components 3 and 8 decline as baboons age. Juvenile baboons contributed more to gray matter covariation components (compared to all other age classes) which include the middle frontal cortex, superior frontal cortex, brainstem, superior parietal cortex, thalamus and pallidum, precentral cortex, insula, and precuneus. In addition, we found adult baboons (of all age classes) contribute more to components 5 and 7 (occipital cortex, thalamus, putamen, and insula). The current findings expand on those reported by Franke et al. (2017) who found reductions in overall gray matter volume as baboons age. Here we show that gray matter covariation within specific regions of the baboon brain change with increased age. As the baboons in the current study had no known illnesses or injuries, the changes in gray matter covariation here can presumably be attributed to normal aging, similar to neurological changes reported in healthy aging humans and chimpanzees (Hopkins, W. D. et al., 2019; Lemaitre et al., 2012; Terribilli et al., 2011). Further, it is possible that these regional changes are related to age-related changes in cognition. In chimpanzees, aged individuals show moderate loss in cognitive and motor functions compared to younger conspecifics (Hopkins et al., 2020a; Hopkins, W.D. et al., 2019; Hopkins et al., 2015; Lacreuse et al., 2018; Lacreuse et al., 2014), and age-related changes in cognition are related to changes in gray matter volume particularly within the frontal and parietal cortices, and precentral cortex (Mulholland et al., 2021). In rhesus macaques, age-related declines in cognitive performance (i.e., delayed response, recognition memory, executive function) are related to decreases in overall volume, synaptic density, and receptor binding density of the dorsolateral prefrontal cortex (Hara et al., 2012). Like humans, chimpanzees, and rhesus macaques, baboon cognition also appears to decline with age; performance on transfer index, novel task learning, precision, and simple discrimination tasks are worse in older adult compared to younger adult baboons (Bonté et al., 2014; Lizarraga et al., 2020); however, additional studies are needed to directly test the relationship between these previously reported cognitive declines and the age-related neuroanatomical differences reported in the current study.

In addition, we found significant sex differences in gray matter covariation. Females contributed more to components which included the brainstem (bilateral), superior parietal cortex (bilateral), thalamus and pallidum (right), precuneus (left), middle and superior frontal cortices (bilateral), amygdala and hippocampus (bilateral), superior temporal cortex (bilateral), and rostral middle frontal (left). Whereas males contributed more to components including caudate and putamen (bilateral), cerebellum (bilateral), and lingual cortex (bilateral). These findings are consistent with other studies which show sex differences in baboon overall brain volume and sulcal length, corpus collosum area, and whole brain gray and white matter volumes (Atkinson et al., 2015; Franke et al., 2017; Phillips and Kochunov, 2011; Rogers et al., 2007). Baboons are sexually dimorphic in both anatomy and behavior. Adult male baboons are larger than adult females, and also display different growth trajectories (with females reaching their adult size at earlier ages than males) and life expectancies (Alberts et al., 2014; Leigh, 2009; Leigh and Cheverud, 1991; Roseman et al., 2010). Sex differences in baboon behavior are also apparent from infancy to adulthood (e.g. communication, social relationships, aggression, play, etc.) (Bentley-Condit, 2003; Brent, 2009; Chalmers, 1980; Coelho and Bramblett, 1981; Pereira, 1988; Smuts, 1985; Young and Bramblett, 1977; Young et al., 1982). As for cognition, there is limited evidence of sex differences to date. Lacreuse et al. (2016) found that males were slower but more accurate on a task of inhibitory control compared to females, perhaps due to testosterone mediated sensitivity to rewards. It is possible that the neuroanatomical sex differences reported in the current study could be related to these sex differences in behavior or cognition; however, more research is needed to examine these relationships.

Though the current study has many strengths, it is not without limitations. The sample was skewed toward younger and female baboons, with half the number of geriatric baboons (n=12) compared to other age classes (juvenile *n*=25, young adult *n*=30, middle-aged adult *n*=22), and almost twice as many females (*n*=58) as males (*n*=31). Although the proportions of males to females reflect proportions within many captive baboon colonies (baboons are typically housed in single- or two-male multi-female groups, i.e. harems), future studies should attempt to include data from more males and ensure that geriatric baboons are represented at numbers similar to other age classes. Despite these limitations, the current study shows that gray matter covariation differs between the two sexes and across age, with geriatric baboons contributing less to covariation components including the frontal and parietal cortices. Future research should examine the relationship between cognitive performance and both age-related and sex-related differences in gray matter covariation.

## Acknowledgements

AM has received funding from the European Research Council under the European Union’s Horizon 2020 research and innovation program grant agreement No 716931 - GESTIMAGE - ERC-2016- STG, from the French “Agence Nationale de le Recherche” ANR-12-PDOC-0014-01 (LangPrimate), ANR-16-CONV-0002 (ILCB) and from the Excellence Initiative of Aix-Marseille University (A*MIDEX). MRI acquisitions were done at the Center IRM-INT (UMR 7289, AMU-CNRS), platform member of France Life Imaging network (grant ANR-11-INBS-0006).

We are very grateful to the Station de Primatologie CNRS, particularly the animal care staff and technicians, Jean-Noël Benoit, Jean-Christophe Marin, Valérie Moulin, Fidji and Richard Francioly, Laurence Boes, Célia Sarradin, Brigitte Rimbaud, Sebastien Guiol, Georges Di Grandi but also the vet Alice Bertello for their critical involvement in this project, Ivan Balansard, Sandrine Melot-Dusseau, Laura Desmis, Frederic Lombardo and Colette Pourpe for additional assistance.

