## Supplemental Materials for "Age- and sex-related differences in baboon (*Papio anubis*) gray matter covariation"

### Slide 1
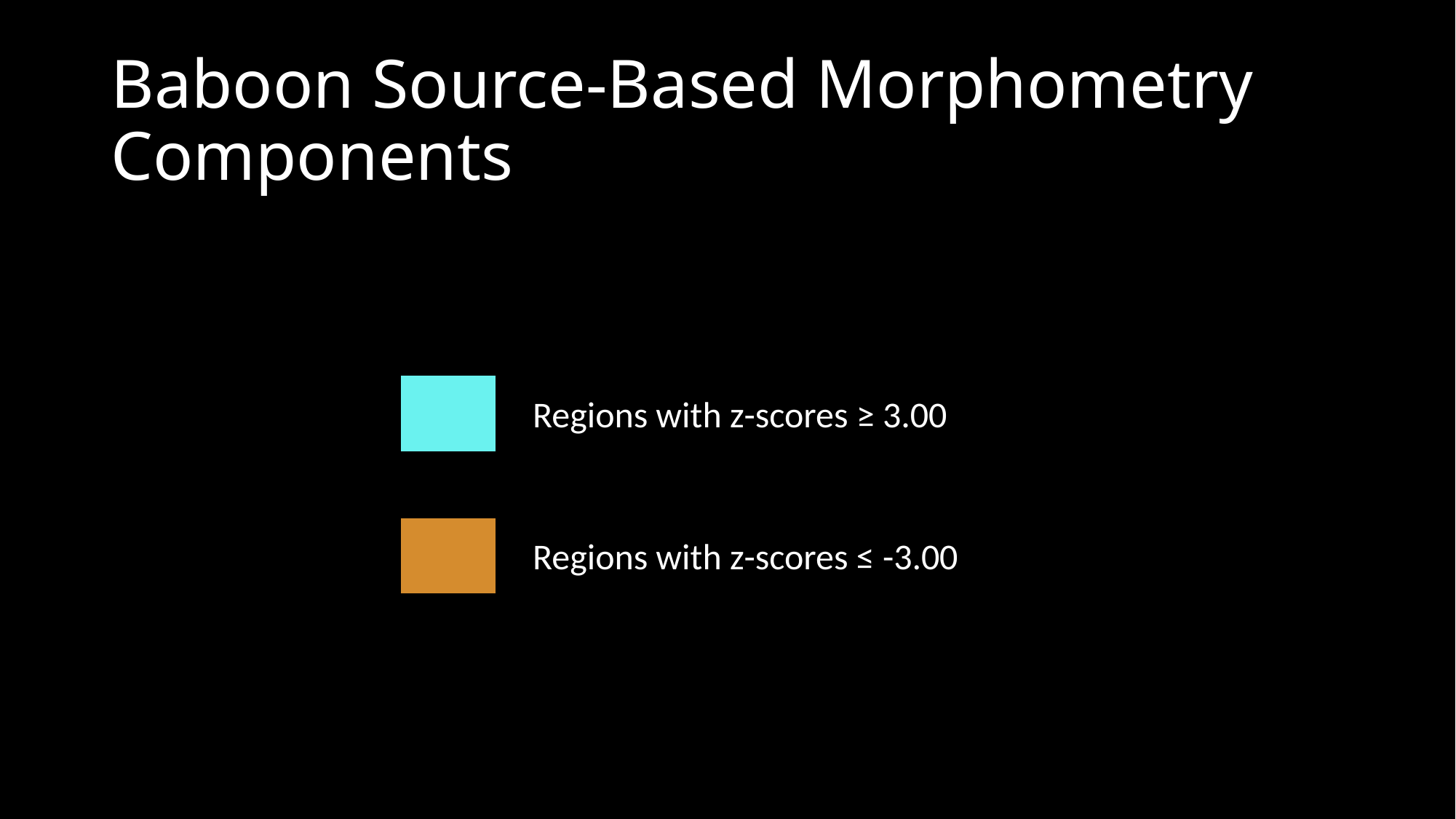

#### Baboon Source-Based Morphometry Components
Regions with z-scores ≥ 3.00
Regions with z-scores ≤ -3.00

### Slide 2
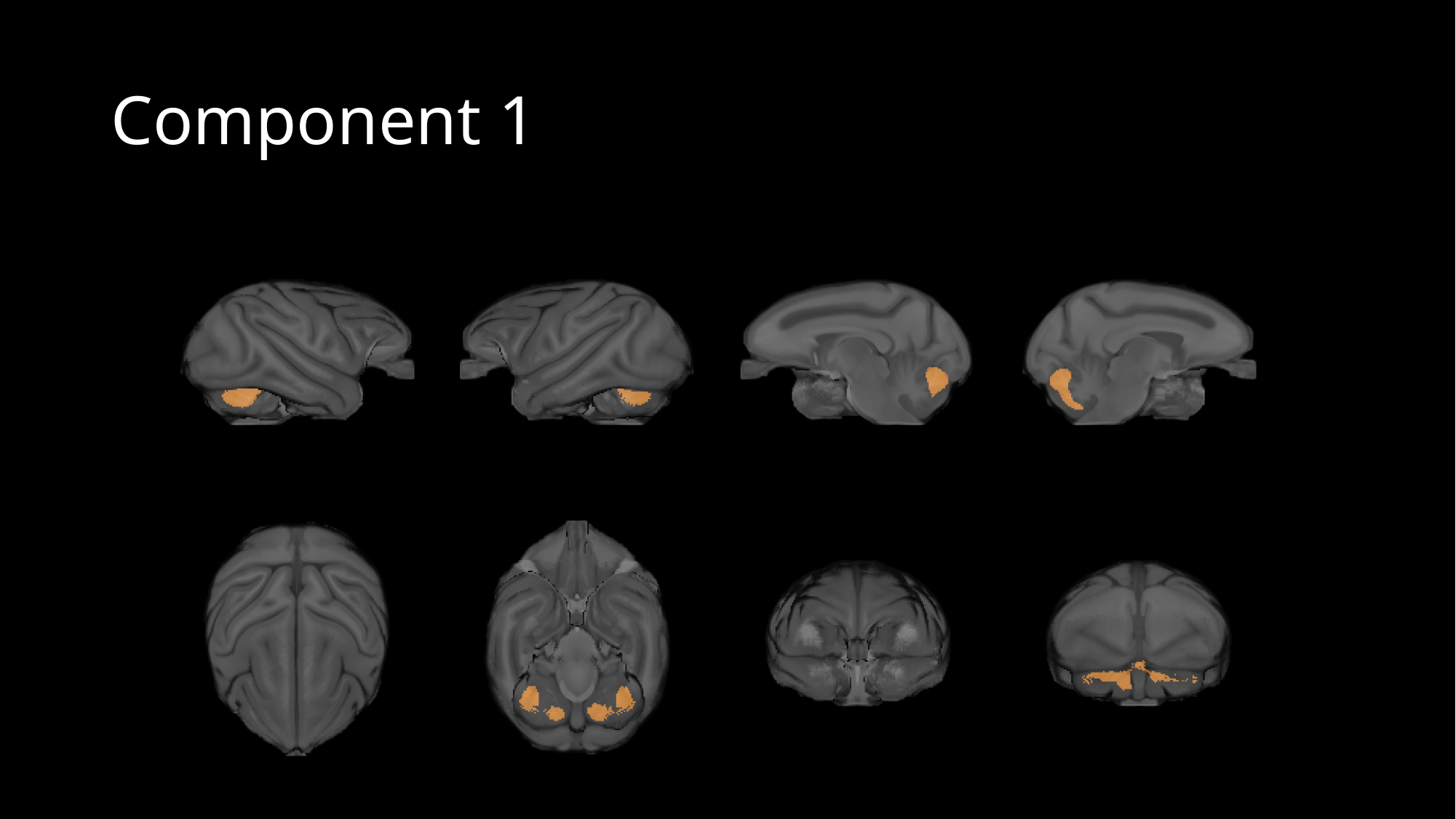

#### Component 1

### Slide 3
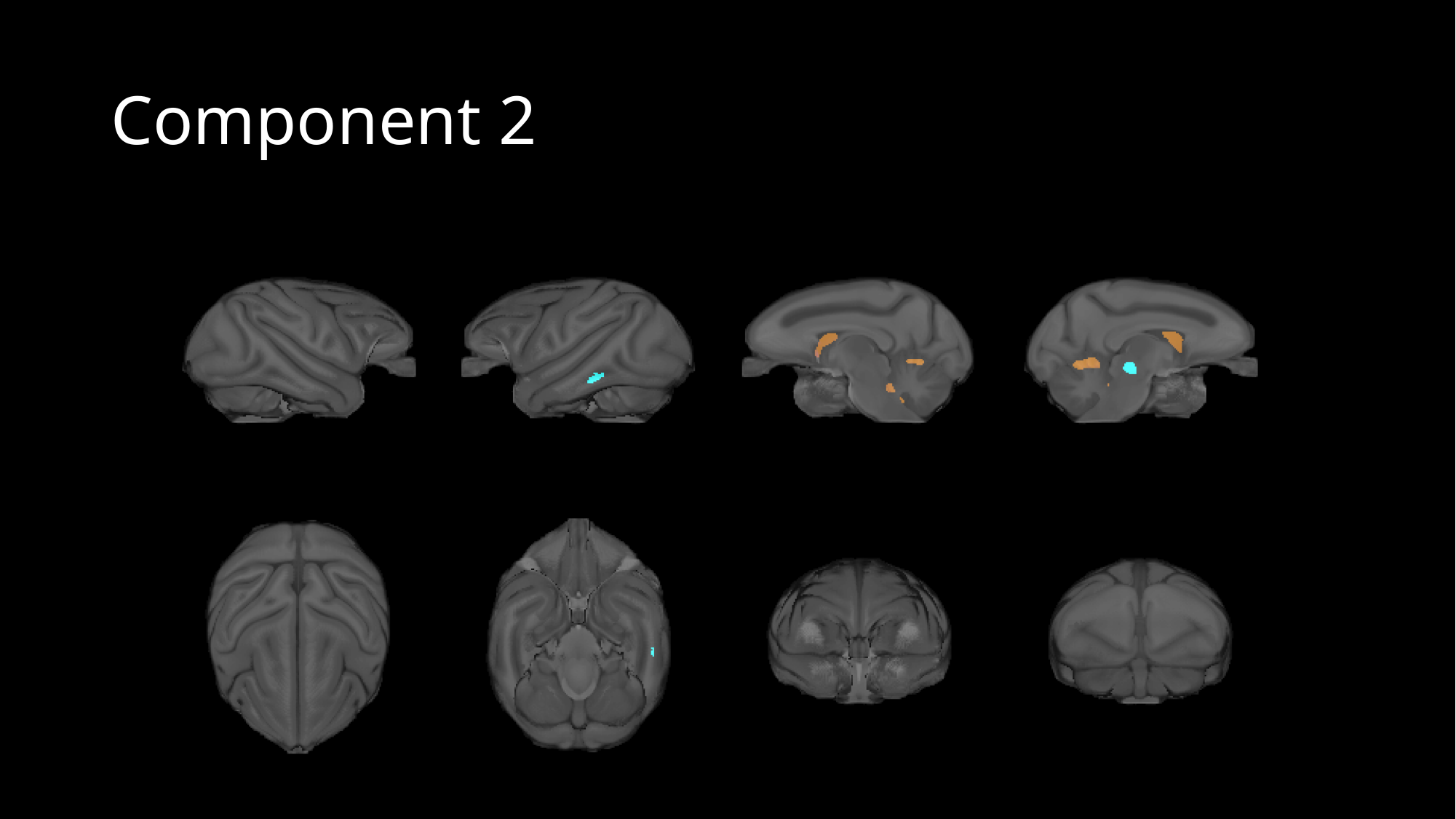

#### Component 2

### Slide 4
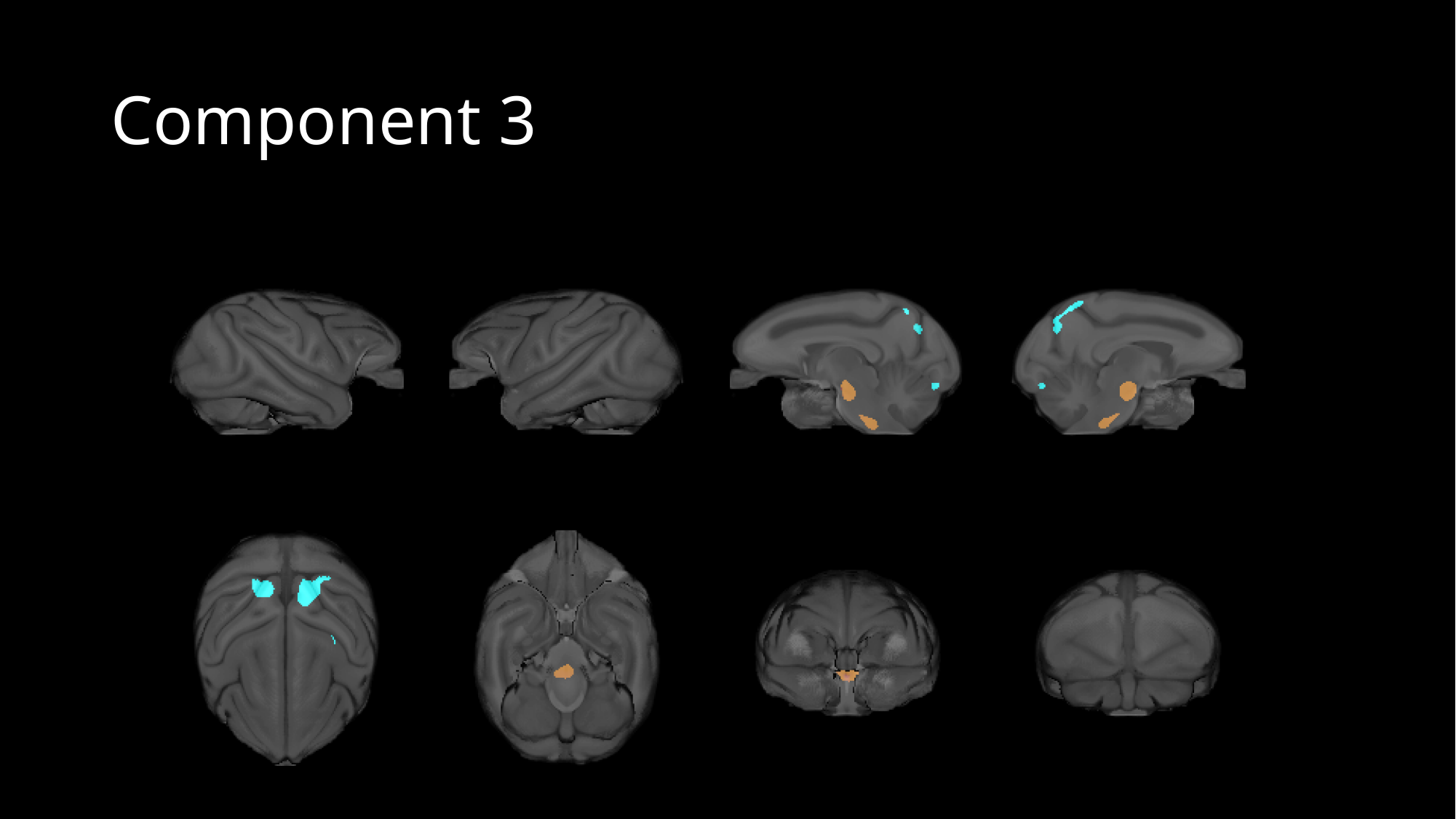

#### Component 3

### Slide 5
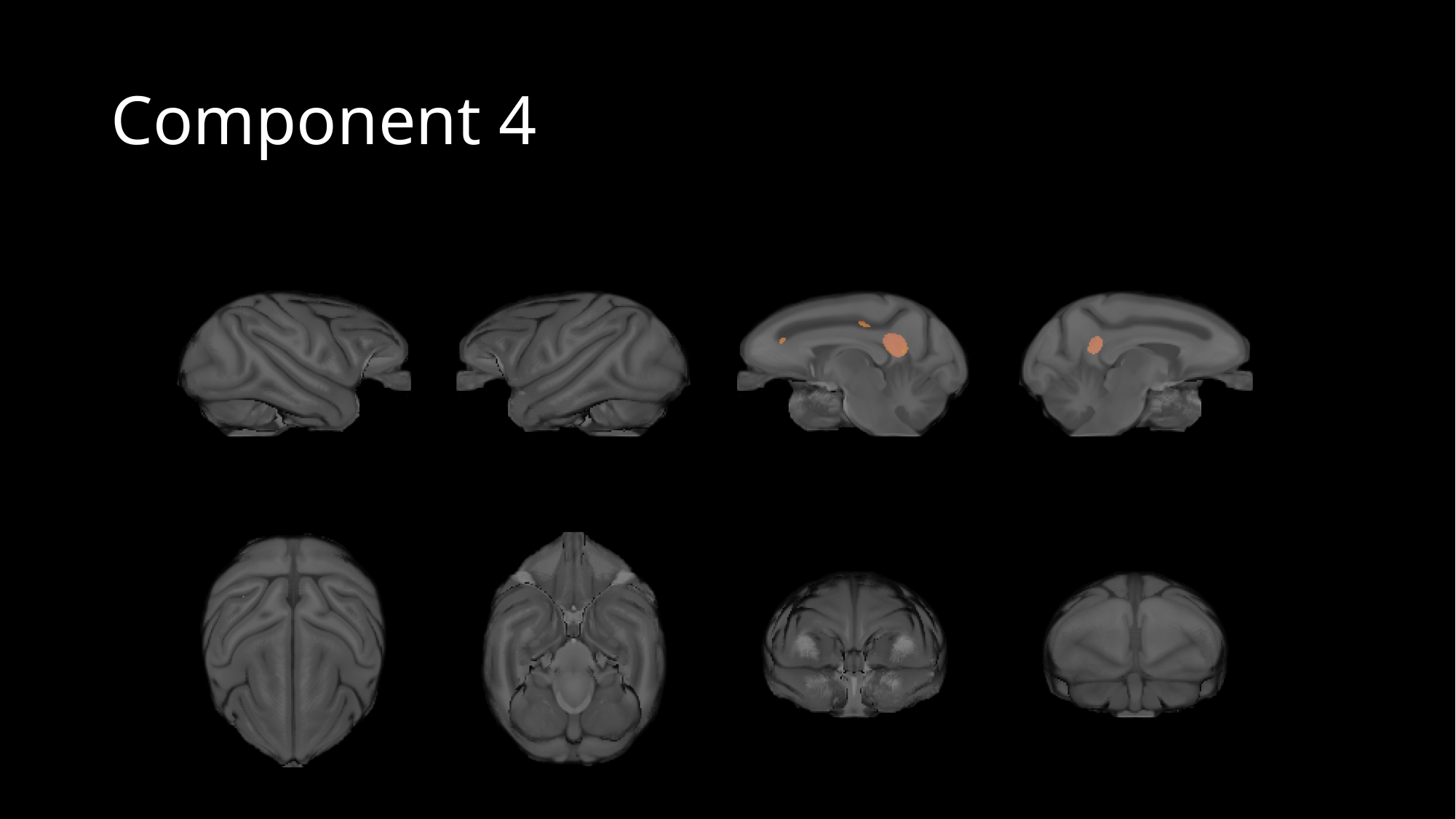

#### Component 4

### Slide 6
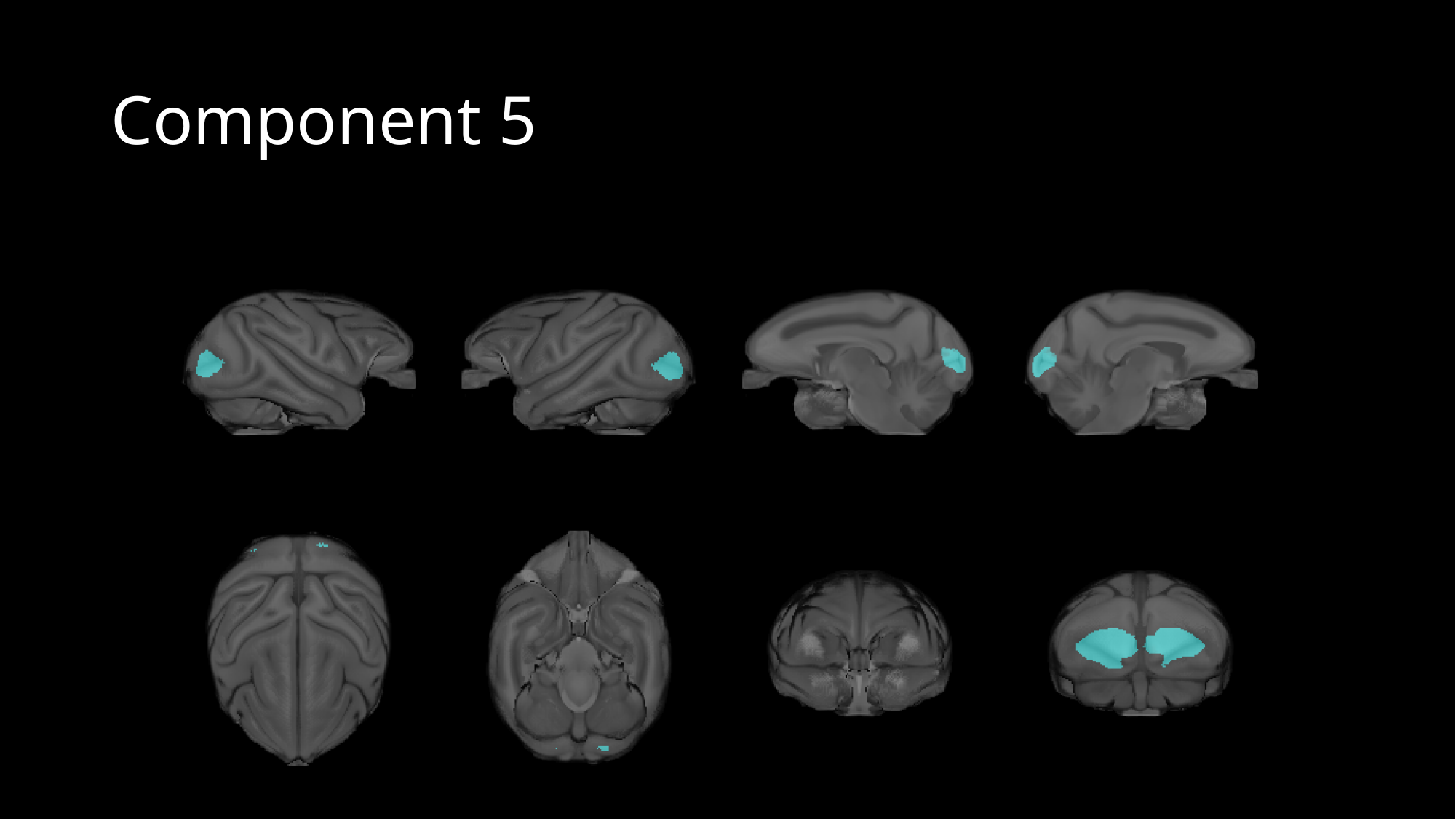

#### Component 5

### Slide 7
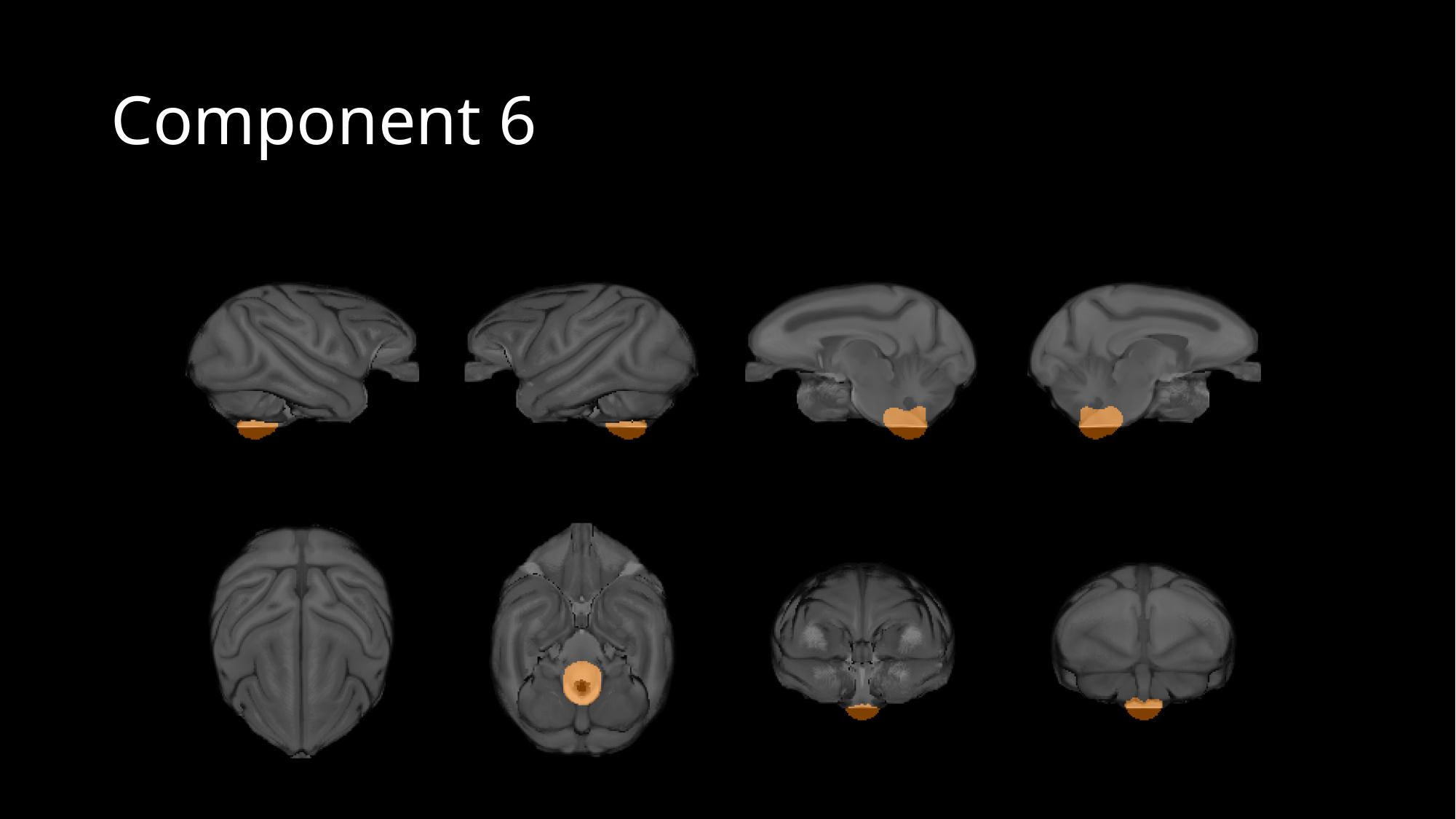

#### Component 6

### Slide 8
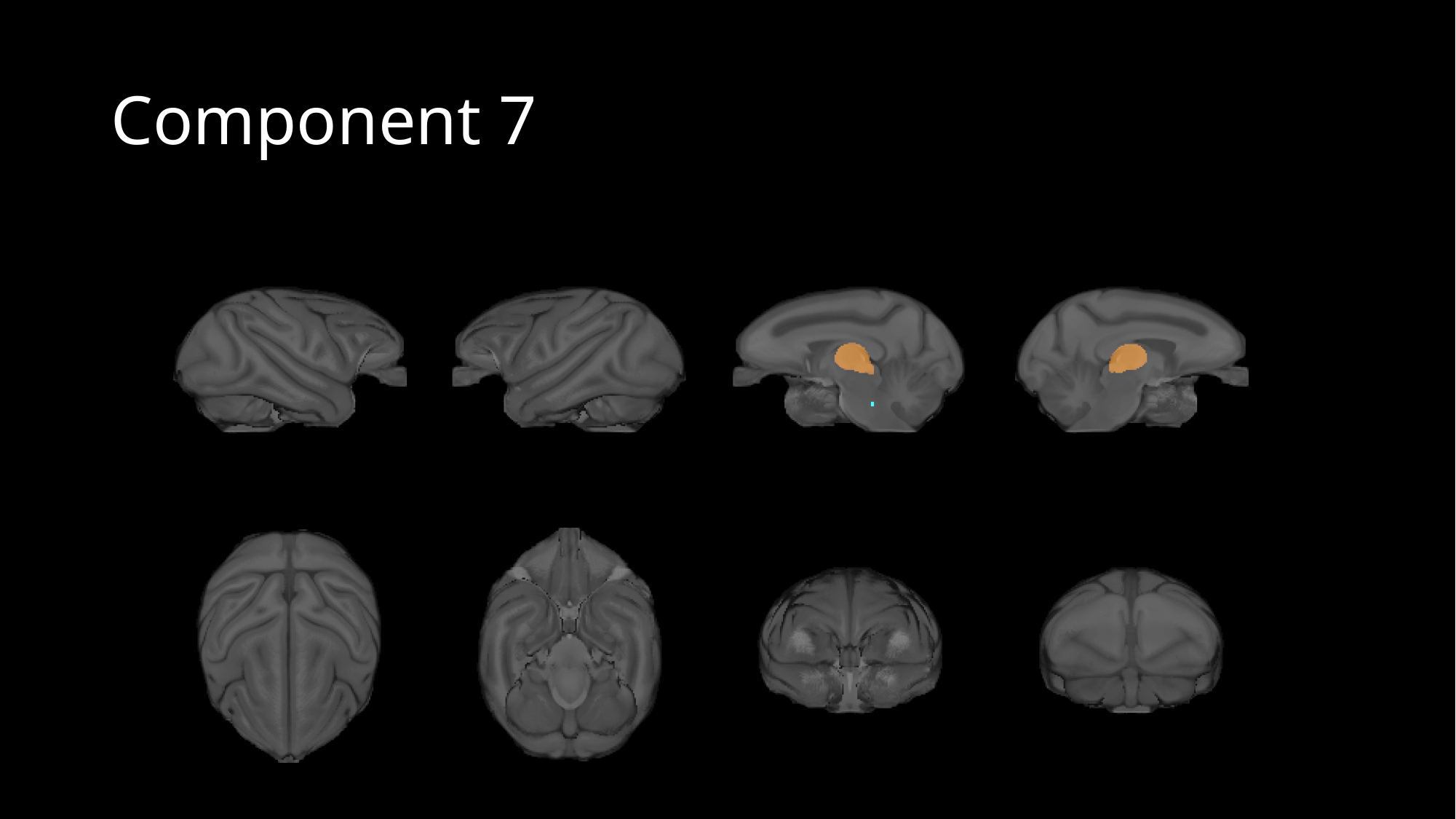

#### Component 7

### Slide 9
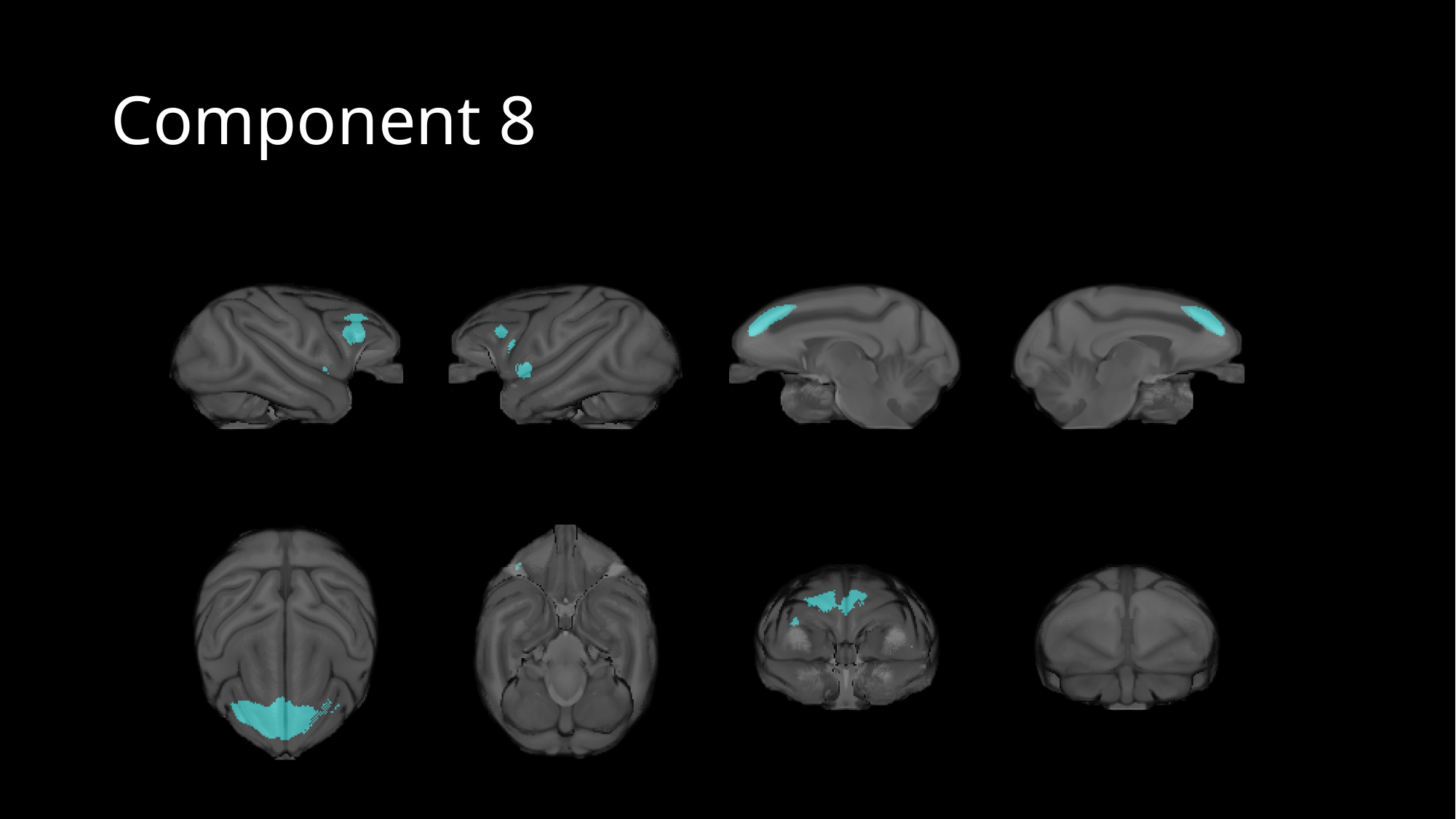

#### Component 8

### Slide 10
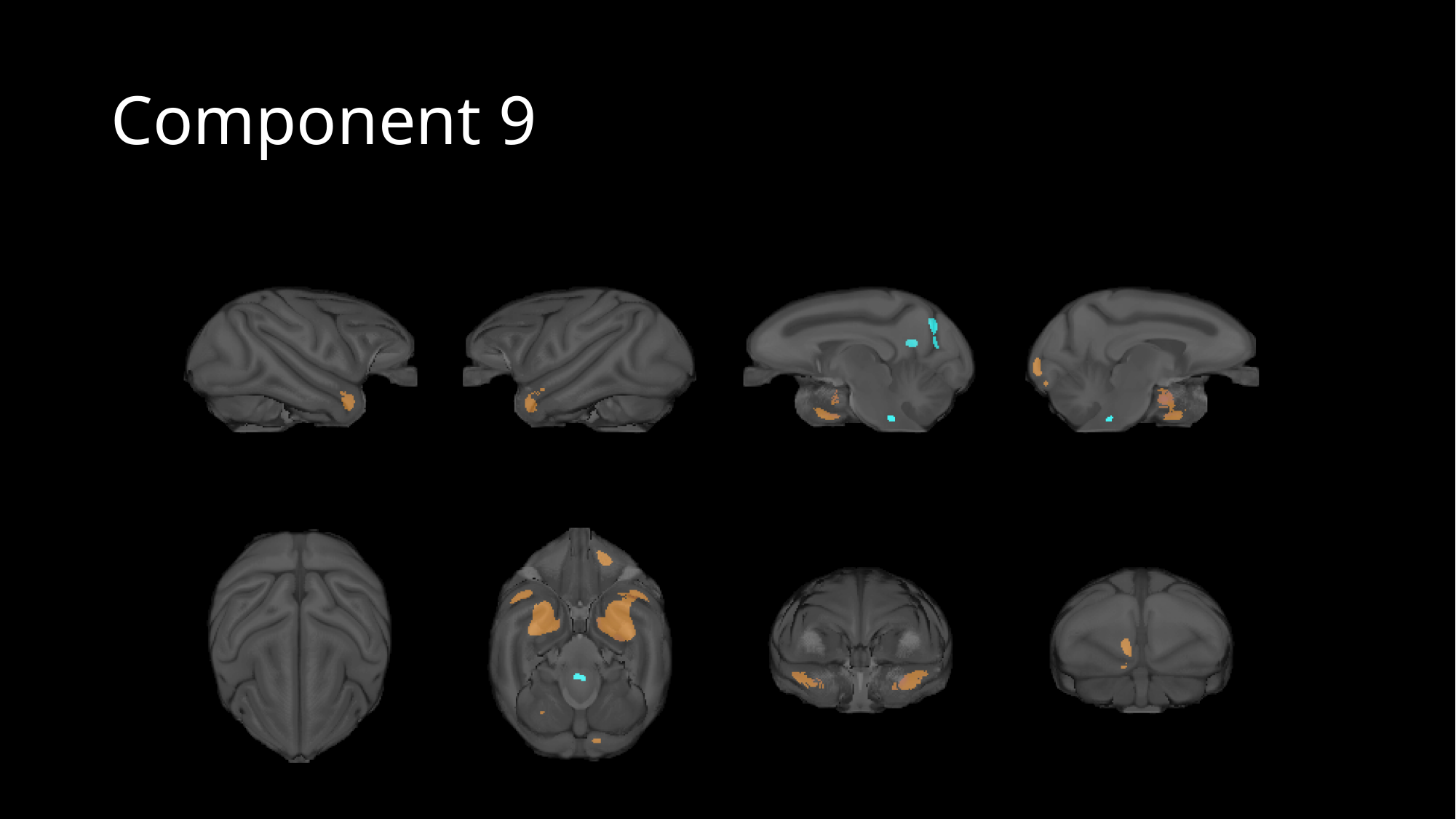

#### Component 9

### Slide 11
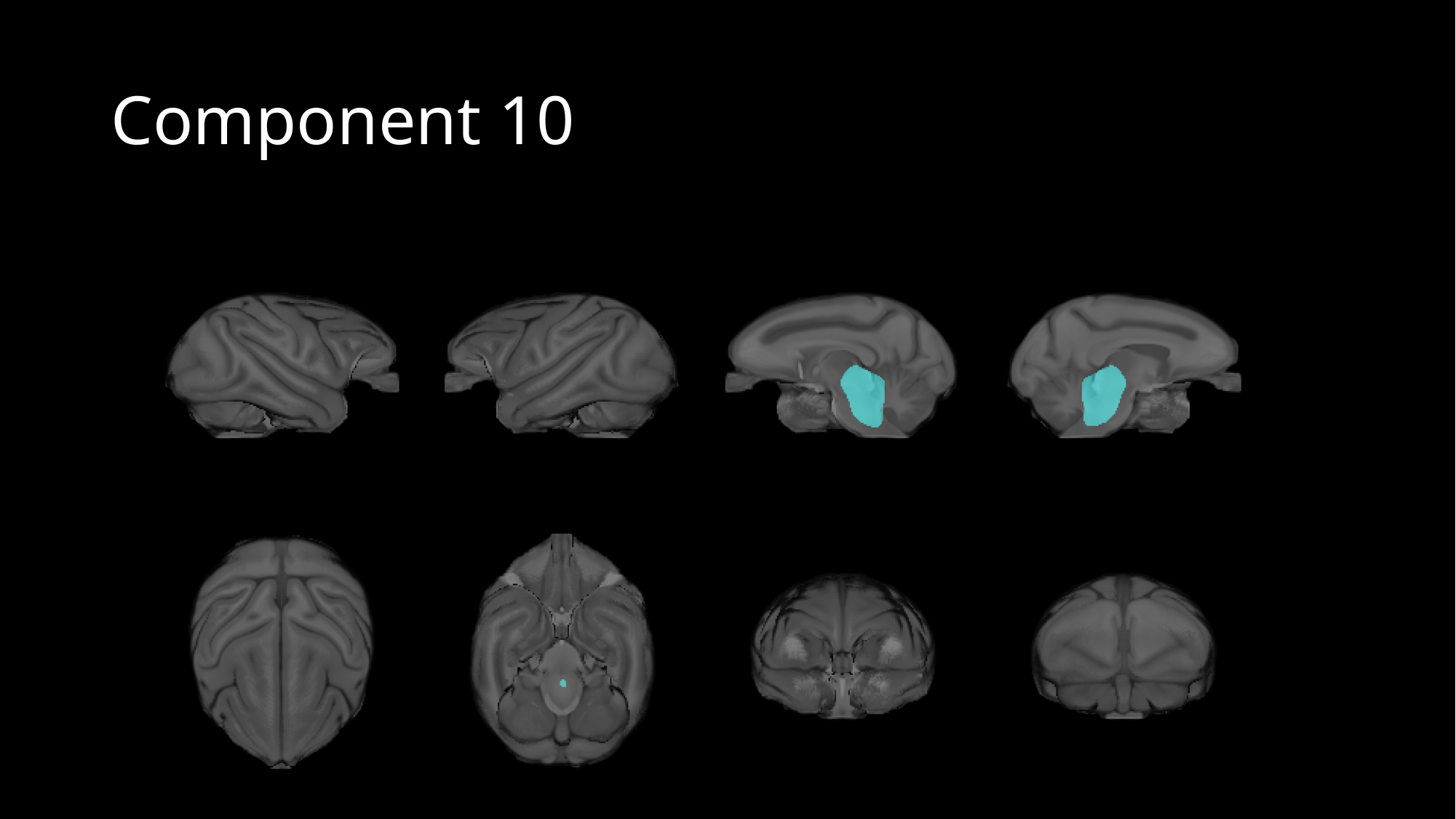

#### Component 10
